# Defining Cell Types through Maximally Informative Biological Atlases

**DOI:** 10.64898/2026.04.23.720468

**Authors:** Roy Wollman

**Affiliations:** University of California, Los Angeles, Department of Integrative Biology and Physiology, Department of Chemistry and Biochemistry, Institute of Quantitative and Computational Biosciences

## Abstract

Cell type atlases organize biological complexity by compressing high-dimensional molecular profiles into discrete cell types that can be mapped across tissues. Yet a principled theory of what constitutes a cell type remains lacking. Here, I introduce an information-theoretic framework in which molecularly defined cell type classifications are evaluated by the information content of the spatial maps they produce. The optimal taxonomy maximizes spatial information, constrained by molecular data, by balancing the entropy of the cell type code against the spatial entropy of the resulting map. The framework generalizes naturally to higher-order spatial partitions, including tissue regions defined by local cellular composition. Applied to a whole-brain mouse spatial transcriptomic dataset, this approach identifies optimal cell-type and region-level taxonomies that balance coding complexity with spatial informativeness. Together, these results establish a unified information-theoretic foundation for cell-type and region-level atlas construction grounded in tissue spatial architecture.

## Introduction

The power of any map, whether geographical or biological, lies in its ability to simplify. Writers such as Lewis Carroll, Jorge Luis Borges, and Umberto Eco famously explored the paradox of a “map the size of the world,” reminding us that models are useful only when they abstract away details [1, 2, 3]. Yet oversimplification is equally futile, leading to the “perfect and absolute blank” of Bellman’s map in Lewis Carroll’s The Hunting of the Snark [4]. This cartographic dilemma is now central to modern biology. The rise of spatial transcriptomics, with its ability to measure gene expression at increasingly high spatial resolution [5], brings this challenge into sharp focus: how to simplify immense biological detail into an appropriate level of abstraction.

Striking the right balance requires a quantitative and objective metric to assess map quality. Information Theory provides the ideal framework for this task, defining a map’s quality through its measurable information content. This approach is well-established in biology, where information-theoretic tools have been used to quantify patterns in gene expression [6], signaling dynamics [7, 8, 9], and their complex dependencies [10]. Most relevant to cartography, pioneering work in the Drosophila embryo estimated that just four genes provide 2.92 bits of spatial information, enough to position a cell with ~1% accuracy [11]. These precedents demonstrate that information theory can provide the rigorous data-driven criterion needed to guide the construction of maximally informative biological atlases.

A central biological instantiation of this abstraction problem is the categorical cell type map. These atlases simplify spatial transcriptomics data by assigning high-dimensional transcriptional profiles to discrete cell type labels, and then mapping those labels onto spatial coordinates. In these types of maps, the cell type taxonomy functions as a legend, a codebook that compresses expression profiles into interpretable units such as “oligodendrocytes.” While such labels do not capture the full richness of the transcriptome [12] or broader aspects of the cellular state [13, 14], they remain valuable abstractions for summarizing spatial transcriptional diversity [15].

Determining what should count as a cell type remains fundamentally unresolved. In practice, taxonomies derived from scRNA-seq are highly sensitive to analytic choices, including the similarity metric, dimensionality reduction, clustering algorithm, resolution parameter, and even the analysis package and software version used [16, 17, 18]. This instability reflects a deeper problem: transcriptomic variation often combines discrete classes with substantial continuous structure rather than forming cleanly separated clusters [19, 20, 21]. The challenge is further complicated by evidence that commonly used dimensionality reduction methods can distort transcriptomic structure and create the appearance of discrete clusters even when the underlying variation is continuous [22, 23]. Even when clusters are present, there remains no principled basis for deciding the appropriate level of granularity. Coarse classifications can erase meaningful distinctions, whereas overly fine ones generate fragmented states with limited interpretability. Molecular data alone therefore do not provide a general basis for deciding which taxonomy should be used.

Even without a principled solution to cell-type definition, the value of biological atlases has led to a rapid expansion of atlas projects. Most have adopted a map-agnostic workflow in which taxonomy is defined from non-spatial data and only then mapped onto tissue. This paradigm became standard in part because single-cell RNA sequencing preceded spatial transcriptomic technologies and offered deeper molecular profiling [24, 25]. Large-scale efforts such as the Human Cell Atlas and the BICCN Brain Atlas exemplify this approach [26, 27]. Yet the instability of purely molecular taxonomies suggests that this approach is unconstrained. Rather than defining a cell-type taxonomy first and then asking where its labels fall in space, a cartographic view treats the taxonomy as the legend of a spatial atlas and asks how well that legend supports the map it produces. Under this view, the challenge is not simply to detect molecular differences, but to determine which categorical partitions, defined by molecular differences, provide the best balance between detail and simplification in spatial organization. The same issue extends beyond cell-type definition. Spatial transcriptomic data can also be partitioned into region types using a growing range of domain-segmentation methods [28, 29], yet these composition-defined maps likewise lack a general criterion for deciding which partition is most informative, and recent benchmarking studies show that no single method performs best across datasets [30]. More broadly, what is missing is a theoretical foundation for defining biological categorical maps, both for cell types and for region types.

Despite these limitations, cell-type taxonomies remain foundational to spatial atlas construction. This study addresses their arbitrariness by providing an alternative map-informed cartographic approach, in which candidate biological partitions, developed from biological features, are evaluated according to the spatial information of the maps they produce. By quantifying the Map Information associated with a given classification, competing cellular taxonomies can be ranked and the one that yields the most informative cellular map can be identified. We further show that Map Information can be extended to anatomical maps in which regions are defined by local cellular composition. We demonstrate this theory by applying it to mouse brain spatial transcriptomics data [27] to identify and characterize information-optimal cell-type and region-level taxonomies.

## Results

### Quantifying the Information Content of Categorical Maps Using Graph Percolation

To quantify the spatial coherence of categorical maps, spatial description can be cast as an address-coding problem. If every cell must be specified individually, the required code length equals the full cellular addressability, *H*_cells_ = log_2_ *N*. Additional information about a cell, such as its cell type, can reduce this code length because different cell types occupy different spatial distributions. A type label is analogous to knowing that an address lies in a residential or industrial area: it does not determine the exact location, but it constrains it. These broad classes can be further subdivided into spatial zones, analogous to neighborhoods within residential areas or industrial parks within industrial areas. Zones therefore provide an intermediate level of description that combines type and spatial information. Rather than specifying each cell directly, the map can be described by first identifying a zone and then specifying position within that zone. Because zones are scale-dependent, their definition depends on the proximity threshold used to construct them. At one extreme, when the threshold is very strict, each cell forms its own zone and the code length is *H*_cells_. At the other, when the threshold is fully relaxed so that all cells of the same type merge into a single zone, the code length reduces to the type entropy, *H*_types_. Evaluating this quantity across a range of spatial proximities captures tissue organization across the scales between these two limits.

#### Box 1

Defining Map Information for Spatial Categorical Maps**x 1: Defining Map Information for Spatial Categorical Maps**

Consider a tissue section with *N* cells, each assigned a type *t* ∈ {1, …, *T*}. The maximal cellular addressability is

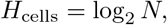

while the non-spatial entropy of the type distribution is

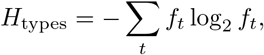

where

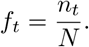

For a proximity scale *P*, let *Ƶ* (*P*) denote the connected components formed by percolation on the type-restricted spatial graph. For two cells *i* and *j* with *t*_*i*_ = *t*_*j*_, we define the edge weight from their rank distance *k*_*ij*_ as

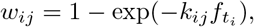

where *k*_*ij*_ is the number of cells expanded to reach *j* from *i* and 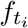 is the fraction of cells of that type. Edges with *w*_*ij*_ ≤ *P* are included at scale *P*. The associated zone entropy is

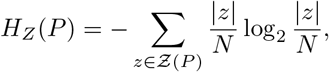

with

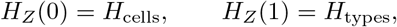

and therefore

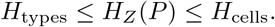

We define Map Information as the reduction in zone entropy produced by the observed spatial arrangement of labels, relative to a label-permuted null, integrated across proximity scales:

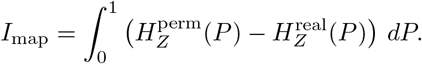

Larger values of *I*_map_ indicate greater spatial coherence.

To operationalize this idea, tissue is represented as a spatial graph whose nodes correspond to cells and whose edges connect cells sharing the same label, with edge weights reflecting spatial proximity (Box 1; Supplementary Theory). For each proximity threshold, only edges below that threshold are retained, and zones correspond to the connected components formed by same-type cells. The resulting entropy curve quantifies how efficiently a labeling compresses cellular addresses through spatial grouping across scales. To isolate the spatial structure contributed by the labeling itself, the observed description length is compared to that obtained after randomly permuting labels across the same set of spatial positions. The difference therefore quantifies how much spatial structure is captured by the classification itself. Classifications that maximize this compression of cellular addresses therefore produce the most informative maps.

Map Information resolves the central tension between maps that are too detailed and maps that are too coarse. If a classification becomes excessively fine, highly specific spatial zones can be obtained, but only at the cost of an increasingly complex code. In the extreme where every cell is assigned its own label, the map retains maximal positional specificity, yet no meaningful compression of cellular addresses is achieved because the type entropy approaches the full cellular addressability. Conversely, if the classification is overly coarse, biologically distinct cells are collapsed into the same label, so the code becomes compact but loses the ability to distinguish important spatial structure. In both extremes, the resulting map fails to provide an efficient description of tissue organization. Crucially, spatial information is used here only as an evaluation criterion: candidate classifications are derived from molecular or phenotypic features independent of space, and their ability to compress cellular addresses is then measured on the spatial map. The most informative atlas is therefore the classification that best balances code complexity with the spatial structure retained by the map.

To illustrate the logic of this framework, we first applied it to a macroscopic test case: the contiguous United States. We constructed three binary county-level maps by dividing counties into two equally abundant classes based on distinct variables: FIPS code parity (even versus odd sum of digits), gross domestic product (above versus below median), and annual average temperature (above versus below median) (Fig. 1a). Because each map was dichotomized at the median, all three labelings have the same type entropy. Differences in *I*_map_ therefore cannot be attributed to class balance or coding capacity, but instead isolate the spatial term in the score: how rapidly the real map percolates relative to its label-permuted null.

**Figure 1.**
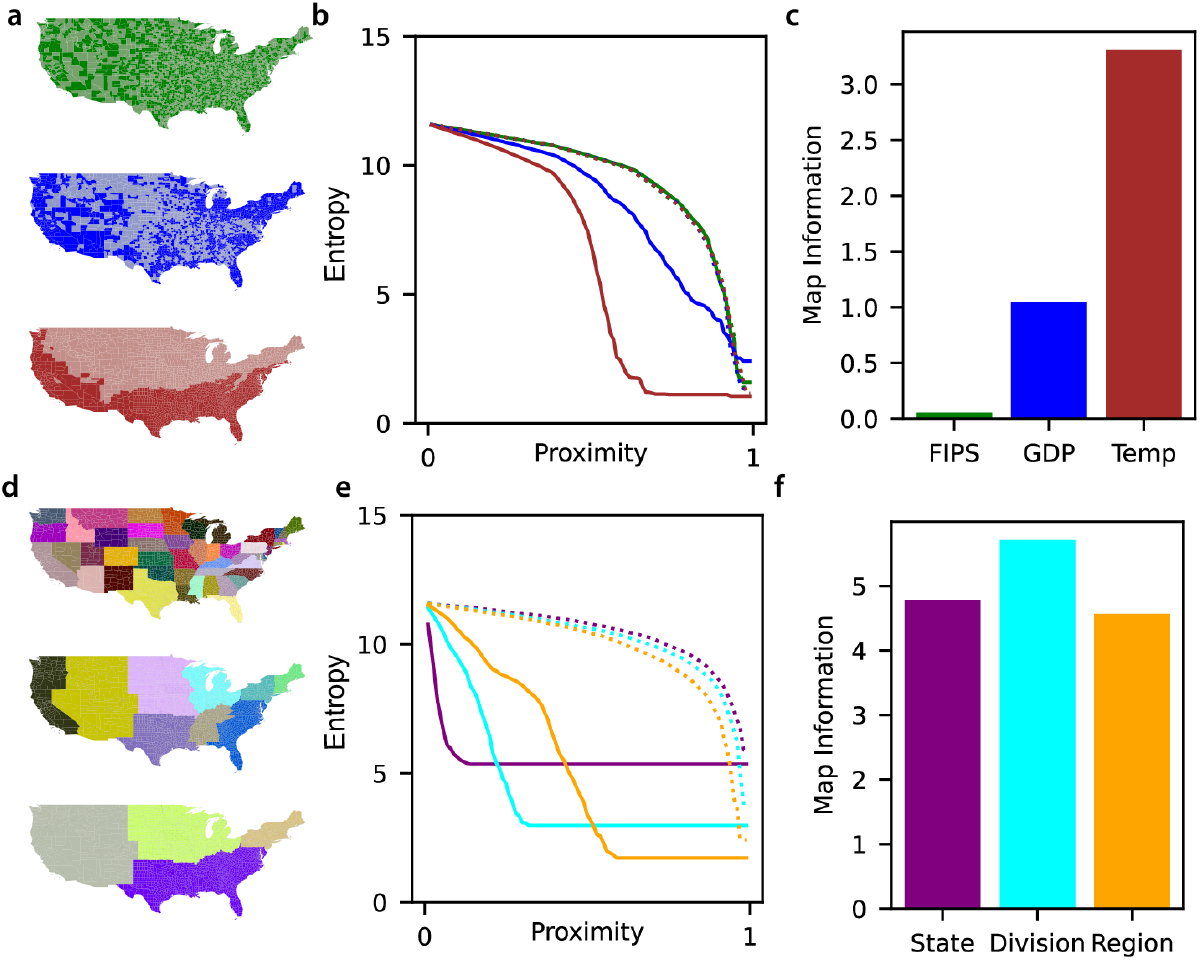
Quantifying spatial information in U.S. maps using percolation analysis. **(a)** U.S. county maps colored according to binary variables with balanced classes (50% abundance each): parity of the FIPS code (green), GDP larger than the median (blue), annual temperature larger than the median (red). **(b)** Percolation analysis of the maps in (a). The curves show entropy as a function of the proximity threshold used to determiine zones, for FIPS (green), GDP (blue), and temperature (red) data. Solid lines are the real data, and dashed lines represent the spatially permuted null models. **(c)** The information score for each map, calculated as the integrated difference between the data and null curves in (b). Temperature exhibits the highest spatial structure.. **(d)** U.S. county maps based on standard U.S. Census Bureau hierarchies: State, Division, and Region, which represent increasing levels of spatial coarseness. **(e)** Percolation curves for the geographic hierarchies in (I): State (purple), Division (cyan), and Region (orange), compared with their respective null models (dotted lines). **(f)** Information scores for the geographic hierarchies. The Division-level classification achieves the highest score, indicating it captures the most significant spatial information by optimally balancing code length and spatial information.

The differences between these maps are visually apparent. The FIPS map appears effectively random in space, the GDP map shows modest clustering around metropolitan regions, and the temperature map forms broad, contiguous latitudinal domains. Percolation analysis captures these distinctions quantitatively (Fig. 1b). For the FIPS map, the zone-entropy trajectory closely follows the spatially permuted null model, indicating that the labeling contributes little spatial organization beyond chance. The GDP map shows a moderate deviation from the null, reflecting localized but incomplete spatial clustering. In contrast, the temperature map exhibits a rapid entropy collapse, consistent with a labeling that partitions the country into large coherent zones. Accordingly, the temperature map achieves the highest Map Information score (Fig. 1c). These examples illustrate the central interpretation of the metric: labelings that create compressible spatial organization produce a larger reduction in zone entropy across scales and therefore higher Map Information.

We then evaluated standard U.S. Census Bureau geographic hierarchies with increasing categorical granularity: Regions (4 categories), Divisions (9), and States (48) (Fig. 1d). This example illustrates the tension between spatial information and type information. The Region map was too coarse to capture substantial geographic structure, whereas the State map required large code to create addresses for all states. The intermediate Division-level classification achieved the highest Map Information score (Fig. 1e,f), indicating the optimal balance between spatial coherence and categorical resolution. Together, these examples show that Map Information does not simply reward spatial contiguity. Instead, it identifies the labeling that most efficiently compresses a spatial map under competing pressures toward oversimplification and over-fragmentation. These examples also motivate the theoretical analysis developed in the Supplementary Theory. There, the metric is shown to have three properties that are directly tested in the analyses below. First, class imbalance limits the maximum attainable score because the available coding range is bounded by type entropy. Second, increasing the number of labels has opposing effects: it can improve spatial coherence by shifting the real percolation curve leftward, but it also reduces the remaining addressable entropy through the coding penalty. This competition predicts a non-monotonic dependence of *I*_map_ on taxonomy granularity. Third, in an unconstrained spatial model the optimum reflects the best possible spatial compression, whereas biological applications are necessarily constrained by molecular or compositional feature spaces. The following analyses therefore ask whether real cell-type and region taxonomies follow these predicted tradeoffs.

### Identifying an optimal cell-type taxonomy of the mouse brain

We then turned to cell-type maps in the mouse brain. As an initial illustration, we examined three representative cell types from the BICCN subclass taxonomy (Fig. 2a). Some subclasses are broadly dispersed across the tissue, whereas others form compact spatial domains. Percolation analysis captures these differences quantitatively. For dispersed cell types, the zone-entropy trajectory remains close to the spatially permuted null model, indicating that the labeling provides little spatial information. In contrast, spatially localized cell types show an earlier entropy collapse and a larger integrated deviation from the null. Thus, even at the level of a single cell type, Map Information quantifies how much coherent spatial structure that type contributes to the atlas.

**Figure 2.**
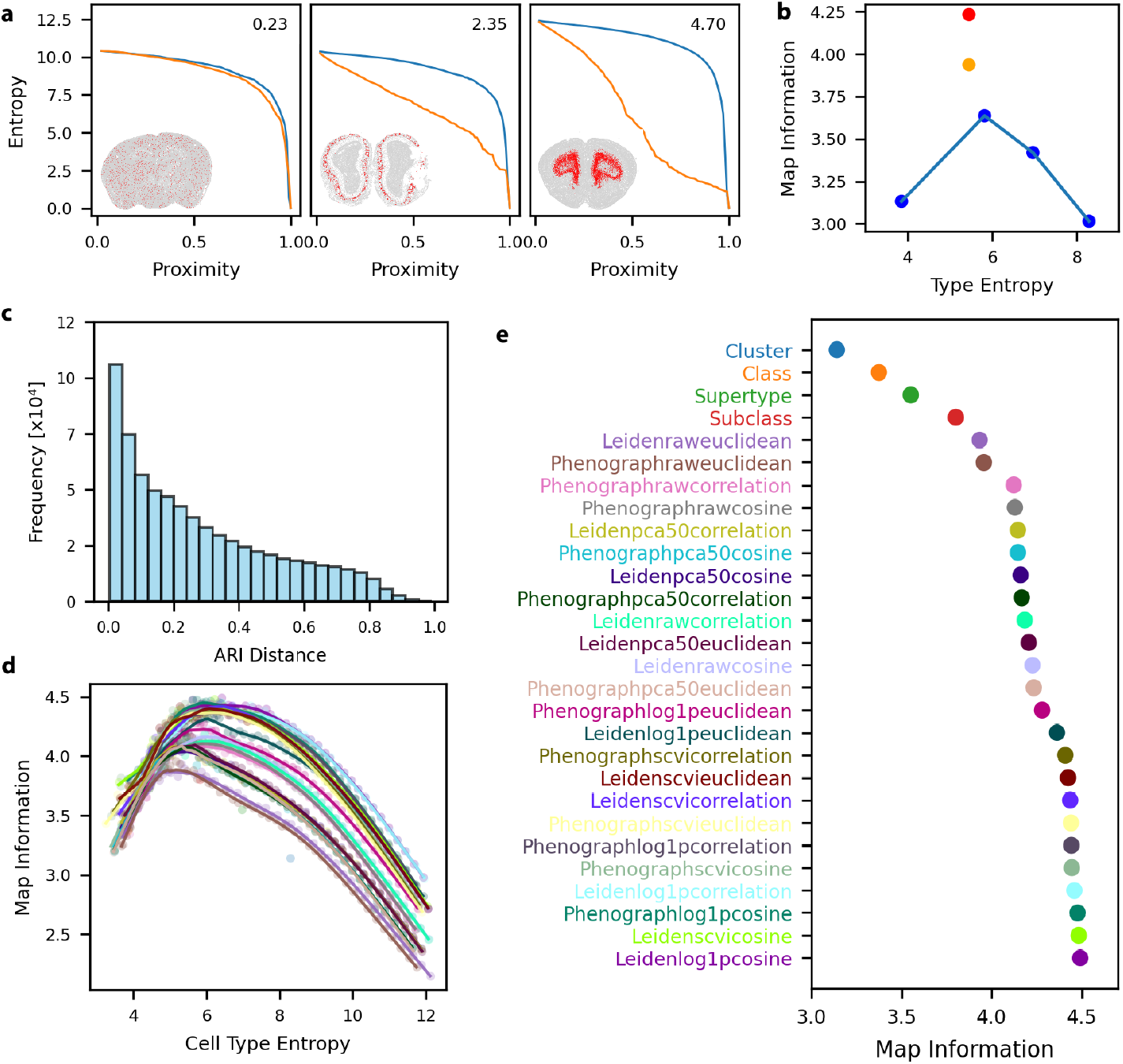
Identification of a maximally informative cell type taxonomy in mouse brain. **(a)**Three examples of single cell type percolation curves left “331 Peri NN”, middle “044 08 Dopa-Gaba” right “030 L6 CT CTX Glut”. The Map Innformation is annotated at the top right corner **(b)** Spatial coherence is plotted against cluster size for the four canonical levels of the Brain Initiative Cell Census Network (BKZCN) taxonomy (blue points). The maximally informative granularities identified by our top-down (red) and bottom-up (orange) heuristic approaches achieve higher coherence than the canonical levels. **(c)** A histogram showing the distribution of pairwise Adjusted Rand Index (ARI) distances between 1,200 unique clustering solutions. The broad distribution demonstrates that the tested algorithms and parameters generated a diverse set of data partitions. **(d)** Map Information as a function of cluster size for all 1,200 tested clustering solutionsu Each point represents a single clustering result, colored by the algorithm used (see legend in e).The plot reveals an optimal range for cluster size that maximizes spatial coherence across different methods. **(e)** Ranking of clustering algorithms by their maximal achieved spatial coherence. This panel provides the color key for the algorithms shown in panel (d).

Theory (see Supp Theory section) predicts that *I*_map_ should be low at both extremes of taxonomic granularity. A very coarse taxonomy has too little label entropy to encode spatial structure, whereas an overly fine taxonomy approaches the cellular-address limit and loses the coding range needed for compression. We tested this prediction by evaluating four levels of the BICCN hierarchy: class, subclass, supertype, and cluster (Fig. 2b). Map Information varied non-monotonically with taxonomic granularity. The class-level map was too coarse to capture substantial spatial structure, whereas the cluster-level map was over-fragmented and incurred a larger entropy penalty. Intermediate levels achieved higher scores, indicating a more informative balance between spatial coherence and categorical resolution. As in the U.S. map example, Map Information identifies the level of abstraction that most efficiently captures spatial organization.

The BICCN hierarchy represents only four possible partitions of transcriptomic space. To test whether nearby alternatives could improve the score, we generated intermediate taxonomies using two simple heuristics: a top-down strategy that progressively split broader groups and a bottom-up strategy that merged finer ones. Both approaches produced taxonomies with higher Map Information than the standard BICCN levels (Fig. 2b), showing that although the subclass level outperforms the other canonical levels, it is not optimal. Yet these heuristic solutions have clear limitations. Because they remain constrained by the structure of the starting hierarchy, they can only partially reorganize the cell-type partition and therefore explore a restricted subset of possible taxonomies.

We therefore expanded the search to a much broader set of candidate taxonomies. We generated 1,200 graph-based clustering solutions of the mouse brain transcriptomic data, organized into 24 families spanning 50 resolutions each. These clustering families differed in data transformation, distance metric, and graph-construction strategy. To assess how distinct these solutions were, we computed pairwise similarity among all 1,200 partitions using the adjusted Rand index. The resulting broad distribution of ARI values confirmed that the clusterings sample a diverse range of transcriptomic partitions (Fig. 2c).

Scoring all 1,200 taxonomies by Map Information revealed a consistent non-monotonic relationship between spatial information and taxonomic granularity across clustering families (Fig. 2d). As the number of cell types increased, Map Information initially rose, reached a maximum, and then declined as further subdivision reduced spatial coherence and increased fragmentation. This behavior matches the theoretical expectation that informative maps must balance gains in taxonomic resolution against the entropy cost of over-partitioning. Importantly, both the height and location of the optimum varied across clustering strategies, indicating that spatial information can distinguish among alternative transcriptomic classifications even when they contain similar numbers of cell types.

Finally, for each of the 24 clustering families, we identified the highest-scoring solution and compared these optima directly (Fig. 2e). The full optimal cell-type atlas is shown in Supp Fig. 1. Several clustering families outperformed both the four BICCN levels and the hierarchy-derived heuristic solutions. Thus, the most spatially informative taxonomy is not recovered by simply selecting one of the standard BICCN annotations or by modestly modifying the reference hierarchy. Instead, it emerges from explicit optimization over a large and diverse set of transcriptomic partitions. Within this framework, the highest-scoring solution provides the cell-type classification that most efficiently organizes spatial structure in the atlas. Importantly, all competing taxonomies evaluated here were derived from molecular features and only then scored by their spatial organization. They therefore differ from classifications defined directly from spatial position alone, which are considered only as an idealized unconstrained model in the Supplementary Theory.

Visual comparison of the BICCN subclass taxonomy and the maximally informative cell-type taxonomy showed that the optimized partition does not broadly scramble reference identities, but instead reorganizes them selectively (Fig. 3a,b). Although it contains fewer categories overall (165 optimized cell types versus 338 subclasses), the optimized taxonomy is substantially more balanced, with a markedly narrower distribution of cells per type (Fig. 3c). In particular, it avoids concentrating nearly half of all cells into only a small number of very large categories.

**Figure 3.**
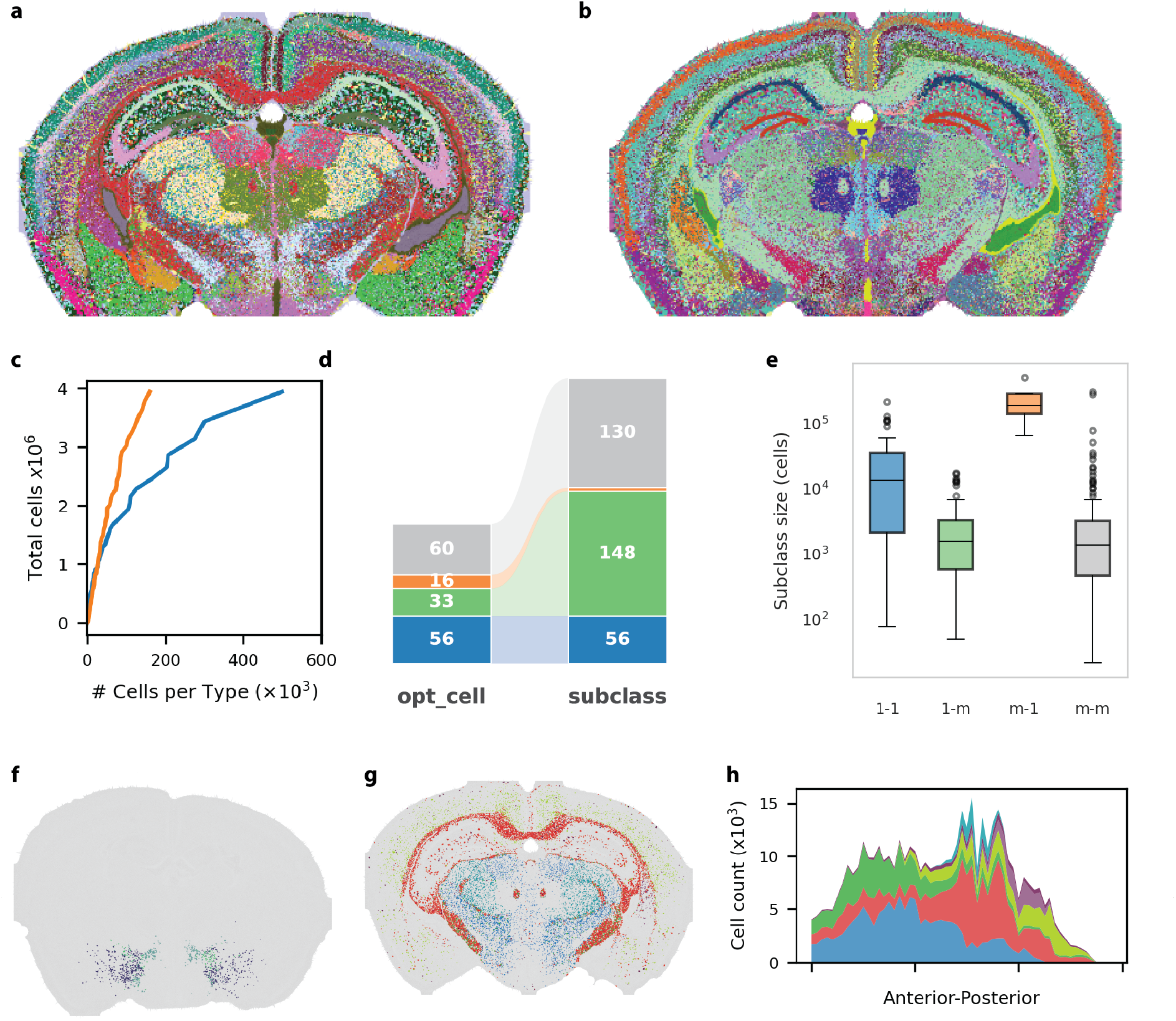
The maximally informative taxonomy redistributes abundant, spatially heterogeneous subclasses into more balanced spatial class. **a,b**, Representative coronal section showed either opt_cell (a) or 8ICCN subclass taxonomies (b). **c** Cumulative distribution of cell-type sizes for the two taxonomies. The optimized taxonomy is more balanced, with a narrower distribution of cells per type despite usng fewer categories (165 optimized cell types versus 338 subclasses). **d** Sankey diagram summarizing type-level correspondence between RCCN subclasses and optimized cell types. Flows are colored by mapping category: one-to-one (blue), one-to-many (green), many-to-one (orange), and many-to-many (gray), highlighting that most relationships remain close to one-to-one. **e** Distribution of the number of cells per subclass across the four correspondence classes shown in d Colors are the same as in d **f** Spatial distribution of the oligodendrocyte subclass in the representative section shown in a **g** The seven optimized cell types that together account for more than 90% of cells assigned to the oligodendrocyte subclass, shown in the same section as f These optimized classes resolve spatial structure that is collapsed within the original subclass annotation. **h** Abundance of the seven optimized oligodendrocyte-derived cell types across the 59 coronal sections. Stacked area plot shows that these dasses form reproducible but dstinct spatial patterns along the anterior-posterior axis.

To quantify the relationship between the two taxonomies, we cross-tabulated cell assignments in both directions and grouped the resulting correspondences into four classes: one-to-one, one-to-many, many-to-one, and many-to-many (Fig. 3d,e; see Methods). The first class was one-to-one, with 56 optimized cell types mapping reciprocally to 56 subclasses, indicating that much of the reference structure is retained. At the same time, 33 optimized cell types represented merges of multiple subclasses, consistent with the overall reduction in category number. In the opposite direction, splitting was concentrated in only four unusually large subclasses, which together gave rise to 16 optimized cell types. Consistent with this pattern, the subclasses involved in one-to-many relationships were substantially larger than those in the other correspondence classes (Fig. 3e). Thus, the optimized taxonomy gains Map Information not by broadly redefining the reference taxonomy, but by merging many small subclasses while selectively splitting a few abundant, spatially heterogeneous ones.

Panels f-h illustrate these two modes of reorganization. Figure 3f shows a representative many-to-one case in which multiple reference subclasses are merged into a single optimized cell type. In contrast, oligodendrocytes provide a clear one-to-many example. In the subclass taxonomy, oligodendrocytes form one broad class, whereas in the optimized taxonomy, this population was partitioned into seven cell types that together accounted for more than 90% of oligodendrocyte assignments (Fig. 3g,h). These optimized classes were not arbitrary subdivisions. In a representative section, they occupied distinct spatial territories, and across the full set of 59 coronal sections they exhibited reproducible but distinct abundance profiles along the anteriorposterior axis (Fig. 3h). Together, these examples show that the optimized taxonomy increases Map Information through selective merging and splitting rather than wholesale reorganization.

### Identifying Optimal Anatomical Taxonomy

We next asked whether the same Map Information framework could be used to evaluate anatomical taxonomies. In the mouse brain, the Common Coordinate Framework (CCF) provides a canonical hierarchy of anatomical annotations spanning five nested levels of resolution [31]. Scoring these five levels showed that the structure level was the most informative among the standard CCF maps (Fig. 4a), showing that the subdivision to substructures reduces labeling efficiency. We then asked whether heuristic refinements of the CCF hierarchy, generated using the same top-down and bottom-up strategy applied in Fig. 2b, could further improve the score. This analysis identified a slightly coarser top-down partition that reached a Map Information score of 8 bits and outperformed the five canonical CCF levels, whereas the bottom-up refinement only improve on the best canonical solution slightly. Thus, within the restricted space of CCF-derived hierarchical refinements, a modestly coarser partition provided a more informative anatomical map than any of the predefined annotation levels.

**Figure 4.**
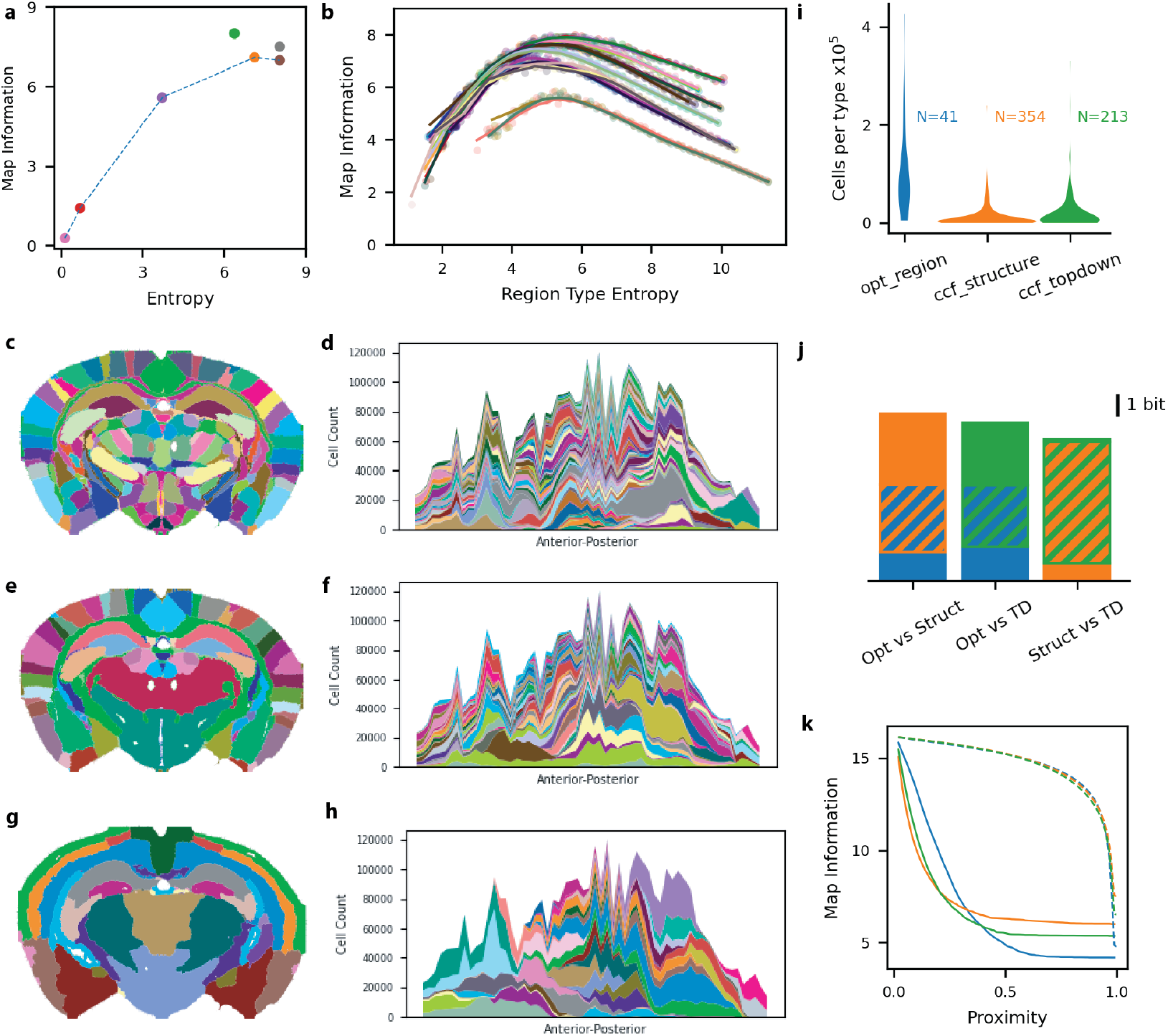
Mop Information identifies informative anatomical taxonomies in the mouse brain. **a**, Mep Wcrmaticn as a function of coding entropy tor the five canonical CCF annotation levels. from organ to substructure. Colored points indicate the five hierarchical levels levels while blue and gray points show the CCF top-down and bottom-up heuristic refinements, respectively. Among the canonical levels, structure achieved the highest Map Information and the CCF top-down refinement further no-eased the score. **b**, Map Information as a function of coding entropy for 1,200 neighborhood-based region partitions generated from local cell-type composition using 24 clustering families spanning 50 resolution each. Faint points show individual clustering solutions and solid curves show smooth fits within each clustering famiy **c,e,g**, Representative coronal sections showing the three region taxonomies compared in this analysis: optimal region (c) CCF structure (e), and CCF top-down (g) **d,f,h**, Abundance of region types across the anterior-posterior axis for the taxonomies shown in c,e,g, respectively. **i**, Distribution of cells per region type for optimal region, CCF structure and CCF top-down. Numbers indicate the total number of region types in each taxonomy. **j**, Entropy decomposition for pairwise comparisons among the three taxonomies. Solid colored segments show conditional entropies unique to each taxonomy, and hatched segments show the shared mutual information **k**, Percolation curves for optimal region, CCF structure, and CCF top-down, with dashed lines shewing the corresponding spatially permuted null models.

Because the CCF was derived largely from classical anatomical criteria rather than local celltype composition, we next asked whether region types defined directly from neighborhood celltype composition could yield a more informative anatomical map. Numerous methods have been proposed to partition spatial datasets into tissue regions based on local cellular composition, underscoring both the biological importance of this level of organization and the lack of consensus on how best to define it [28, 29, 30]. As with transcriptomic clustering, the resulting partitions depend strongly on analytic choices, including neighborhood definition, similarity metric, dimensionality reduction, and resolution [12, 17, 18]. We therefore used Map Information as an external evaluation criterion to compare alternative neighborhood-based partitions and to identify a maximally informative region taxonomy for the mouse brain.

To search broadly for such a taxonomy, we generated 1,200 neighborhood-based clustering solutions for the 4 million cells in the mouse brain using local cell-type composition, with neighborhoods defined from the maximally informative cell-type taxonomy identified in Fig. 2. These solutions were organized into 24 clustering families spanning 50 resolutions each, varying in neighborhood definition, dimensionality reduction, and similarity metric. Pairwise adjusted Rand index comparisons confirmed that these solutions sampled a broad and diverse set of region partitions. Scoring all 1,200 solutions by Map Information revealed a consistent non-monotonic relationship between Map Information and regional coding entropy across clustering families (Fig. 4b). When the partition was too coarse, the taxonomy failed to capture the full spatial complexity of the tissue, whereas overly fine partitions incurred a penalty for excessive code complexity. Across clustering strategies, intermediate values of regional coding entropy produced the highest Map Information scores. Comparing the top-scoring solution from each family then identified the optimal region taxonomy.

The optimal region taxonomy achieved a higher Map Information score than CCF structure and matched the CCF top-down partition, with both reaching 8 bits. The full view of the optimal region type categorical atlas is shown in (Supp. Figure 1). We therefore compared these three anatomical maps directly (Fig. 4c-h). Although CCF top-down and optimal region achieved similar scores, they did so using markedly different partitioning strategies. Visually, the optimal region taxonomy used labels more economically, a pattern confirmed quantitatively by the distribution of cells across region types (Fig. 4i). The optimal region taxonomy contained only 41 region types, each represented by a relatively large number of cells, whereas CCF structure used 354 types, many of which were sparsely populated. CCF top-down was more parsimonious than CCF structure, with 213 region types, but remained substantially more complex than the optimal region solution. Consistent with their common origin, CCF top-down and CCF structure showed high mutual information with each other, whereas their mutual information with the optimal region taxonomy was more modest relative to the entropy of each partition (Fig. 4j). These differences were also reflected in the percolation curves: compared with CCF top-down, the optimal region taxonomy collapsed more slowly at short range but reached lower entropy at larger spatial scales, indicating that similar overall Map Information can arise from distinct tradeoffs between local fragmentation and large-scale spatial compression.

### Cross-level information mapping

After identifying the maximally informative cell-type and region taxonomies, we asked how strongly these labels were specified by the feature spaces from which they were derived. Using a 500-gene panel, we defined 165 cell types, and using local cell-type composition, we defined 41 region types. We therefore quantified the mutual information between each taxonomy and subsets of its underlying features. As the size of randomly sampled gene subsets increased, so did the information they provided about cell-type identity, such that the full 500-gene panel recovered nearly all of the information contained in the cell-type labels, with mutual information approaching the entropy of the taxonomy itself (Fig. 5a). Thus, the selected cell-type labels are tightly constrained by transcriptional space, with nearly all of their coding entropy recoverable from the full 500-gene panel. An analogous analysis of region labels as a function of neighborhood cell-type features showed the same overall behavior (Fig. 5b): the full feature set captured nearly all of the information in the regional taxonomy, whereas progressive feature removal reduced the recoverable information. Together, these results indicate that both taxonomies are strongly grounded in the molecular or compositional feature spaces from which they were derived, rather than reflecting arbitrary categorical assignments.

**Figure 5.**
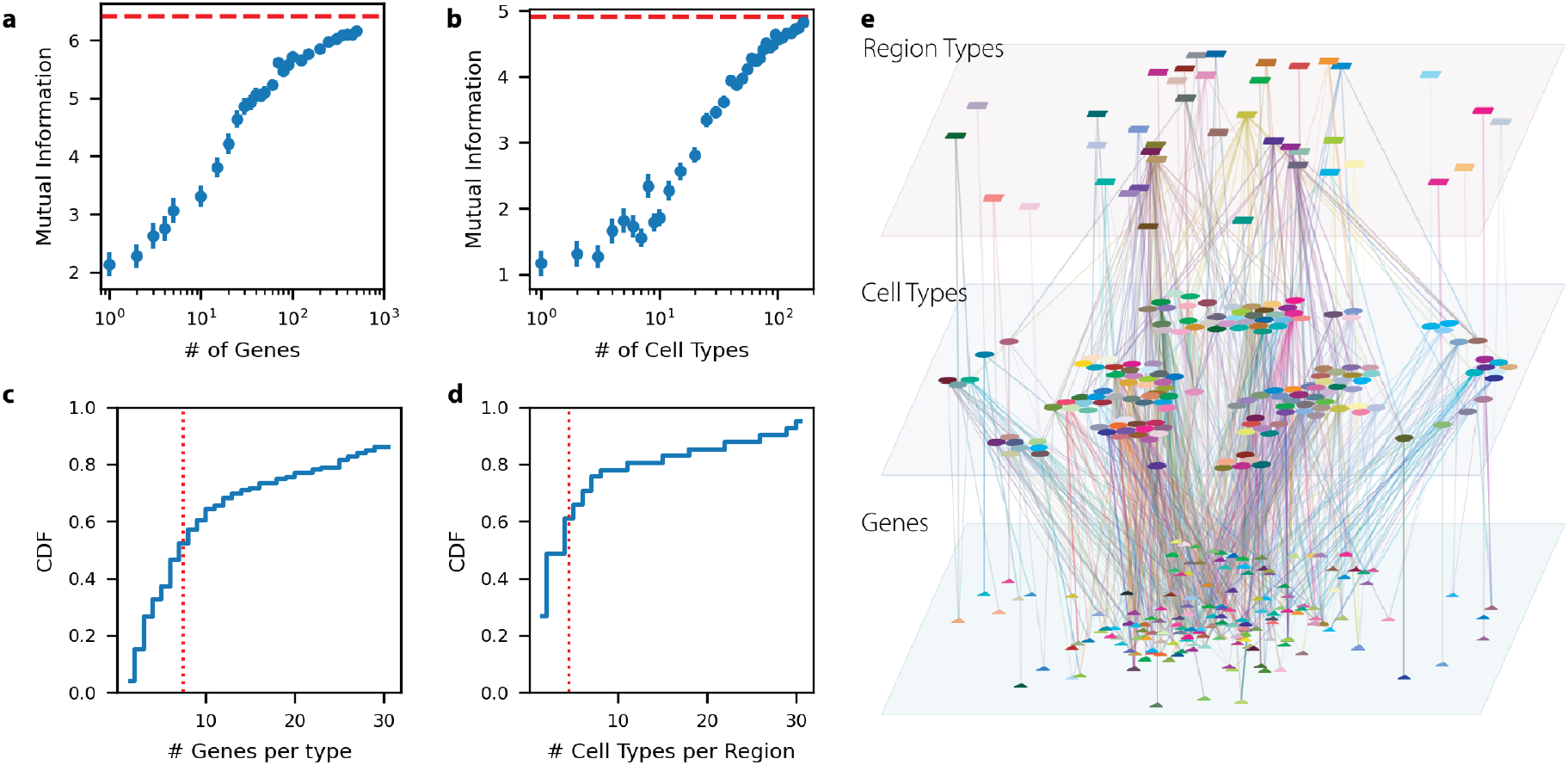
Genes and local cell-type composition define cell-type and region-type taxonomies through sparse, partially overlapping feature sets. **a**, Mutual information between randomly sampled subsets of the 500-gene panel and the 165-cell type taxonomy. Points show mean ± SEM across sampled gene sets. As additional genes are included, recoverable information approaches the entropy of the full cell type taxonomy (red dashed line). **b**, Mutual information between randomly sampled subsets of neighborhood cell-type features and the 41-region-type taxonomy. Points show mean ± SEM across sampled feature sets. Increasing the number of cell-type features similarly increases recoverable information toward the entropy of the full regional taxonomy (red dashed line). **c**, Cumulative distribution of the number of genes required to reach a mutual information of OS for individual cell-type labels using a greedy feature-selection procedure Red dashed line marks the median. **d**, Cumulative distribution of the number of cell-type features required to reach a mutual information of 08 for individual region-type labels, analyzed as in c Red dashed line marks the median. **e**, Multi-level graph linking the 256 genes selected in at least one greedy feature set to all 165 cell types and all 41 region types. Cell-type and region-type node colors match the taxonomies shown elsewhere in the paper, whereas gene colors are used only to distinguish nodes. Edges summarize feature usage across greedy feature sets, with more prominent edges indicating more frequent selection.

We next asked how many features were required to specify individual labels within each taxonomy. For each cell type or region type, we constructed a balanced one-versus-rest classification problem so that the maximal label entropy was 1 bit, and then used a greedy search to identify the smallest set of genes or cell-type features that maximized mutual information with that label. Individual labels could often be recovered with relatively sparse feature sets, although the number required varied substantially across labels. The median number of genes needed to reach a mutual information of 0.8 bits for cell-type labels was 8, whereas the analogous median for region labels was 4 cell-type features (Fig. 5c,d). Rather than being specified by entirely unique marker sets, individual cell types and region types were typically defined by relatively small combinations of features drawn from a partially overlapping repertoire. Some labels could be identified by one or a few highly specific genes or cell-type features, whereas others required larger combinatorial signatures, indicating that the taxonomy is organized through sparse but reused building blocks rather than isolated one-to-one markers. We summarized this sparse and partially overlapping organization as a multi-level graph linking genes to cell types and cell types to region types (Fig. 5e).

## Discussion

From the earliest days of cell biology to modern single-cell genomics, what constitutes a cell type has remained unresolved. Despite the centrality of this question, the field has lacked a theoretical foundation for determining cell-type taxonomy, resulting in subjective determination and a large number of opinion pieces [32, 33, 34, 35]. In this study, I introduce an objective framework based on information theory for evaluating cell-type taxonomies through the maps they produce. The framework formalizes the cartographic principle that a useful map must balance detail against simplification [36]. It treats cell-type taxonomy as a map legend and finds the optimal taxonomy as the one that best represents tissue organization with the most efficient legend. The core contribution of this work is the introduction of a new information theory metric, Map Information, that formalizes this idea by treating a biological map as a problem of spatial address compression, in which the taxonomy serves as the categorical code for describing cellular organization across scales. Under this view, the mere existence of molecular differences is not enough to define a cell type, avoiding the subjective challenge of determining how many molecular differences are enough to create a new cell type. Cell types are instead justified by whether the resulting taxonomy yields an informative spatial compression of tissue organization. Applied to a whole-brain mouse spatial transcriptomic dataset, this framework identified information-optimal cell-type and region-level taxonomies, illustrating how Map Information can provide a general criterion for evaluating biological partitions in practice.

A unique feature of this framework is that it distinguishes the problem of defining cell types from the statistical problem of finding structure in molecular feature space. Conventional clustering criteria such as silhouette width, likelihood-based penalties, or information criteria can evaluate compactness, separation, or model fit, but they do not address the biological question of which molecular distinctions should be elevated into cell-type boundaries [12]. In practice, because cell states often lack inherent separability [19], different statistical criteria rarely converge on the same solution, making the choice among them inherently subjective. Map Information addresses a different problem. Rather than asking whether a partition is statistically tidy, it asks whether that partition yields an informative map of tissue organization. In this sense, it provides an external criterion for evaluating candidate taxonomies, one grounded not in the internal geometry of transcriptional space alone, but in the spatial compression that the resulting labels achieve in tissue.

The mouse brain analyses illustrate several consequences of using a theoretically rigorous framework. First, the most informative classifications emerged at intermediate levels of granularity rather than at either extreme, consistent with the underlying cartographic tradeoff between excessive detail and excessive simplification. Second, the optimized cell-type taxonomy did not arise through wholesale reorganization of the reference hierarchy, but through selective merging and splitting. Many small subclasses were effectively collapsed, whereas a smaller number of broad and spatially heterogeneous groups were partitioned into more informative subtypes. Third, the same general logic extended beyond cell types to anatomical organization: both canonical CCF partitions and neighborhood-derived region taxonomies could be evaluated under the same criterion, revealing that distinct anatomical schemes can achieve similar overall information values while doing so through different tradeoffs between local fragmentation and large-scale spatial compression. Fourth, the resulting multilayer classification followed a few-to-few mapping across the hierarchy from genes to cell types to region types. Together, these results suggest that Map Information is not merely a way to score clustering outputs, but a framework for comparing alternative biological partitions across levels of organization.

At the same time, information-optimal does not necessarily mean universally adopted or practically dominant. Biological taxonomies, like other coding systems, are shaped not only by formal efficiency but also by history, convention, and the need for shared standards. The QWERTY keyboard is a familiar example: not the most efficient arrangement, but highly functional because of its widespread adoption [37] The same is likely true for reference atlases such as the BICCN hierarchy [38, 39] or the CCF [31]. Even when alternative partitions preserve more spatial information, established taxonomies retain major value because they support comparison across studies, accumulation of annotations, and community interoperability. Map Information should therefore not be viewed as a mandate to replace existing standards, but as a principled framework for quantifying their tradeoffs, for identifying when alternative classifications may be more informative for specific atlas-building goals, and for shaping the design of future atlas projects, particularly in nascent systems where standards have not yet solidified.

The current framework is also limited by the space of candidate solutions supplied to it. Map Information is an evaluation criterion, not a discovery engine in itself, and the optima identified here should therefore be interpreted as the best solutions recovered within a substantial but still restricted search over candidate partitions. Although the present study explored 1,200 diverse classification options, this search remains only a partial sampling of the much larger space of possible molecular and compositional classifications. A more systematic strategy for navigating partition space, ideally one designed explicitly around the optimization target rather than inherited from standard clustering workflows, will be needed to determine how close these solutions are to a true global optimum. Developing such a strategy remains an open problem, because Map Information is sensitive to spatial organization but not to molecular detail per se, making it nontrivial to use directly as an optimization target without drifting toward spatially coherent but molecularly unconvincing classifications.

A second limitation concerns the data itself. The present analysis is based on two-dimensional tissue sections, whereas the underlying biological problem is inherently three-dimensional at the scale of the whole organ. In addition, the quality of the inferred taxonomies depends on the nature and accuracy of the molecular measurements. The optima identified here should therefore be viewed as conditioned on the data provided. As data quality improves and whole-organ three-dimensional profiling matures, the resulting classifications and their associated Map Information values will likely improve as well. In this sense, the objective criterion provided by Map Information offers an additional benefit: it may help indicate when improvements in experimental data collection have become sufficient, as reflected by convergence in Map Information.

Overall, Map Information provides a theoretical foundation for defining biological taxonomies that are spatial in nature. A deeper question is whether the compressed spatial address favored by Map Information reflects something biologically meaningful beyond descriptive parsimony. One possibility is that these optima simply provide the most efficient summaries of data generated by processes organized around entirely different constraints. Another is that biological systems themselves evolved in ways that make such compressed spatial descriptions informative for development, physiology, or function [13]. Whether this reflects a deeper principle of biological organization, perhaps because evolution, morphogenesis, or physiological demands favor tissue architectures that are efficiently describable across scales, remains open.

### Code availability

All custom code and software used for the computational workflows and statistical analyses in this study are open-source and publicly available. The core tissue multigraph analysis and visualization package (TMG) can be accessed on GitHub at https://github.com/rwollman/TMG. The `max info atlas` pipeline, which provides modular feature extraction, clustering, percolation analysis, result aggregation, and high-performance computing (HPC) orchestration, is available at https://github.com/rwollman/max_info_atlas.

## Methods

### Data sources

All analyses were performed on a whole-brain mouse spatial transcriptomic dataset stored as an AnnData object (`C57BL6J-638850-raw.h5ad`) provided by [27]. Anatomical annotations were derived from the Allen Institute CCF annotation matrix (`annotation 10.nii`). Analysis of the US map in Figure 1 was done based on Bureau of Labor Statistics (FIPS score: https://www.bls.gov/cew/classifications/areas/qcew-area-titles.htm), Bureau of Economic Activity (GDP data: https://www.bea.gov/data/gdp/gdp-by-county), and National Oceanic and Atmospheric Administration (Temperature: https://www.ncei.noaa.gov/access/monitoring/climate-at-a-glance/county/mapping)

### Spatial graph construction

For each tissue section, cells were represented as nodes in a spatial graph defined by their two-dimensional tissue coordinates. Rather than using Euclidean distance directly in the percolation calculation, spatial proximity was encoded through rank distance. For a given cell *i*, all other cells were ordered by spatial proximity, and the rank *k*_*ij*_ of cell *j* in this ordering was used as the fundamental distance measure. This rank-based formulation reduces sensitivity to local density variation and makes the graph construction more comparable across tissue contexts with different sampling densities.

For a fixed labeling, percolation was evaluated on the subgraphs induced by cells sharing the same label. Thus, the relevant graph at each step was not a general cell-cell graph with arbitrary edges, but a same-label proximity graph whose connectivity depended on both spatial rank and label frequency. For two cells *i* and *j* with label *t*_*i*_ = *t*_*j*_, the edge strength was defined as

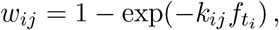

where 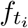 is the global frequency of label *t*_*i*_. This frequency-dependent normalization was introduced so that a fixed threshold would correspond to a comparable notion of spatial enrichment for rare and common labels. Rank distance was used rather than Euclidean distance in order to reduce distortions caused by spatial density heterogeneity across tissue architecture. Rank distances were calculated with exact KNN using SciPy implementation of KNN search when the search included *<* 100, 000 cells and using pynndescent [40] for cases with *>* 100, 000 cells.

### Percolation analysis and Map Information computation

To calculate Map Information, we performed a deterministic bond percolation analysis on type-restricted spatial graphs. Initial local connectivity was established using a *k*-nearest-neighbor algorithm, where potential edges were strictly limited to nodes of the same type. Each edge was assigned a deterministic activation threshold (*p*_bond_) proportional to its continuous neighbor rank and the global frequency of that specific node type. Because local nearest-neighbor graphs fail to capture tissue-wide connectivity at the limit of high *k*, we introduced a macroscopic bridging approximation. This algorithm temporarily condenses locally connected cells into isolated zones, computes their geometric centroids, and calculates a rank-based distance matrix to establish physically symmetric, macroscopic edges between distant islands of the same type. This macroscopic bridging step was used only to approximate large-scale same-label connectivity in the finite local *k*-nearest-neighbor graph and does not change the definition of Map Information in Box 1, which is based on the entropy of connected components in the thresholded type-restricted graph.

Once all local and macroscopic edges were aggregated and sorted by their *p*_bond_ thresholds, we performed a percolation sweep from 0 to 1. The edge-merging process was accelerated using a custom Cython-compiled union-find data structure optimized with path compression and union-by-size. Rather than recalculating the entire graph state at each threshold, the algorithm maintained a dynamic dictionary cache of component-size frequencies, allowing for *O*(1) updates during merges. To further minimize the computational overhead of floating-point operations, the Shannon entropy of the connected components was calculated at approximately 1% intervals of the total edge list.

For each tissue section, the section-level Map Information score was computed as the integrated difference between the entropy curve of the randomly permuted null model and that of the observed spatial graph,

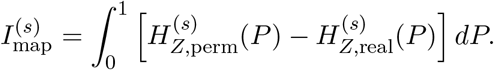

The integral over *P* was approximated numerically from these sampled entropy values using the trapezoidal rule over the corresponding *p*_bond_ thresholds, with *P* = 0 and *P* = 1 included explicitly.

The null entropy curve was generated by randomly permuting labels across the fixed cell coordinates within that section, thereby preserving the section-specific label counts while destroying spatial organization. The same section-wise permutation procedure was used for cell-type, region-type, and CCF-derived maps.

For whole-brain analyses, Map Information was computed independently for each coronal section and then averaged across sections as a cell-weighted expected value,

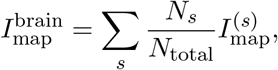

where 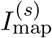 is the score for section *s, N*_*s*_ is the number of cells in that section, and *N* _total_ = ∑_*s*_ *N*_*s*_.

### Hierarchy-derived merge and split heuristics

To maximize spatial informativeness, we applied top-down (splitting) and bottom-up (merging) greedy heuristics to refine the reference cell-type hierarchy. Both approaches explored intermediate taxonomies constrained by established parent-child relationships. To ensure computational efficiency, the total Map Information score was formulated as the sum of independent, pre-computed contributions from each selected cell type. A type’s contribution was calculated by weighting its spatial entropy difference curve by its global frequency, summing across tissue sections, and integrating over the probability threshold.

The top-down algorithm initialized at the broadest hierarchical level, iteratively replacing parent nodes with their direct subtypes if the substitution strictly increased the total score. Conversely, the bottom-up algorithm began at the finest leaf-node resolution, merging sibling subtypes into their immediate parent to improve the score. These data-driven refinements allowed the taxonomy to autonomously adapt to the optimal spatial resolution supported by the underlying tissue architecture.

### Generation of candidate transcriptomic clustering families

A broad search over alternative transcriptomic partitions was done by creating many variants of the core Leiden clustering algorithm [41]. The variants used four feature spaces (`raw`, `log1p`, `pca50`, `scvi`), three graph distances (`correlation`, `cosine`, `euclidean`), two graph-clustering methods (`leiden`, phenograph [42]), and 50 resolution settings for each family. This yields 4 × 3 × 2 = 24 clustering families and 24 × 50 = 1, 200 candidate transcriptomic clustering solutions. A 15-nearest-neighbor graph was used in this sweep. The diversity of the set of 1,200 classification was verified by calculating all pairwise Adjusted Rand Index (ARI) values.

### Comparison between optimized and reference cell taxonomies

To compare the optimized cell taxonomy with the reference subclass taxonomy, we cross-tabulated cell assignments and established significant correspondence using a reciprocal 66% cell mass coverage threshold. Specifically, for each optimized cluster and reference subclass, we identified the minimal set of partner categories required to encompass at least 66% of its assigned cells. Based on the cardinality and mutual exclusivity of these reciprocal sets, we classified the cluster relationships into four topological categories: one-to-one (mutually exclusive single partners, indicating conserved cell types), one-to-many (one optimized cluster merging multiple reference subclasses), many-to-one (multiple optimized clusters splitting a single reference subclass), and many-to-many (complex, non-exclusive overlaps). Finally, the distributions of cell counts per type were compared across these topological classes to assess the relative granularity and scaling of the optimized taxonomy.

### Generation and evaluation of candidate region taxonomies

Region taxonomies were generated by clustering local cellular composition feature matrices using a pipeline analogous to the cell-type optimization. First, the optimal cell-type annotations (`LeidenLog1pCosine_res2p683`) identified in Figure 2 were used to define local microenvironments. For each cell, spatial neighborhoods were determined using *k*-nearest spatial neighbors at multiple scales (*k* ∈ {10, 25, 50, 100}). Unweighted raw counts of neighboring cell types were computed to create four base local frequency feature matrices. To capture major axes of compositional variation, principal component analysis (PCA, *n* = 15 components) was applied to each scaled base matrix, yielding four additional derived feature matrices, resulting in a total of eight feature families.

From these eight feature representations, *k*-nearest neighbor graphs (*k* = 15) were constructed using cosine, Euclidean, and correlation distance metrics, producing 24 distinct graph topologies. Candidate region taxonomies were then generated by applying the Leiden community detection algorithm to each graph across 50 log-spaced resolution parameters (ranging from 10^−2.0^ to 10^1.5^). To enforce spatial contiguity, a spatial smoothing step was applied post-clustering using a Delaunay triangulation adjacency graph, which merged disconnected small components (fewer than 50 cells) into their surrounding regions. This comprehensive parameter sweep generated an exact total of 1,200 candidate region taxonomies (8 feature families × 3 distance metrics × 50 resolutions), which were subsequently evaluated using spatial percolation analysis to identify the optimal regional scale.

### Mutual information analysis linking genes, cell types, and region types

To quantify the degree to which specific feature sets specify the optimized taxonomies, we employed an information-theoretic framework. We calculated the mutual information (MI) linking continuous gene expression profiles to discrete cell types, as well as local cellular composition features to region types. To robustly estimate MI between continuous and categorical variables, we utilized a *k*-nearest neighbor (KNN) approach [8, 43]. An approximate KNN graph was constructed for the continuous feature space using nearest neighbor descent with a correlation distance metric. Because empirical MI estimators are subject to sample-size-dependent bias, we applied an asymptotic bias correction. MI values were calculated across varying computational subsample sizes (*N*), and the bias-corrected true MI was derived by fitting the relationship to the asymptotic model MI(*N*) = *A* − *B* · *N*^−*C*^, where the asymptote *A* represents the corrected MI estimate.

To identify compact feature subsets sufficient for reconstructing the target labels, we implemented a greedy forward-selection procedure. To ensure computational efficiency and balanced class representation during this search, the cellular space was subsampled to 2,000 cells per iteration, comprising 1,000 cells from the target classification and 1,000 randomly selected cells representing all other types. Starting with the single most informative feature, additional features were iteratively added by maximizing the joint mutual information between the expanding feature subset and the target labels. To ensure numerical stability and prevent distance collisions during high-dimensional KNN computations, continuous features were standardized and a uniform micro-jitter was applied. This selection process continued up to a predefined maximum feature limit, ensuring the selection of a highly informative, compact alphabet. The robustness of these selected feature trajectories was evaluated by calculating the standard error of the mean (SEM) of the information gain across 10 independent sampling repeats.

Finally, these relationships were conceptualized as a tripartite graph connecting genes, cell types, and region types. Edge weights in this network were defined by the selection stability of each feature during the optimal greedy search. Specifically, for each target classification, we identified the minimum number of selection steps required to reach a predefined mutual information threshold (0.8 bits). We then evaluated all independent sampling repeats that achieved this threshold in the minimum number of steps. The edge weight between a feature and a target type represents the fraction of these optimal repeats in which the feature was selected, yielding a stability score between 0 and 1. This provided a robust, systems-level visualization of how consistently specific, spatially resolved transcriptional features drive local tissue organization.

## Supplementary Text: Characterization of the Map Information metric

This Supplementary Text has two distinct purposes. First, it describes the implemented rank-distance, same-label spatial graph used to compute Map Information. Second, it develops simplified analytical models that provide intuition for the behavior of the metric. These analytical models are intentionally idealized. They are not intended to reproduce the full geometry, finite-size behavior, graph construction, or biological constraints of the empirical analyses. Instead, they isolate specific effects: the normalization of rank-distance edges, the tradeoff between coding complexity and spatial coherence, the effect of class imbalance, the contrast between feature-constrained and unconstrained spatial partitions, and the expected behavior under simple random label noise.

### 1 The Construction of the Spatial Graph

The implemented Map Information calculation represents each tissue section as a spatial graph whose nodes are cells and whose candidate edges connect cells assigned to the same label. Edge weights are based on rank distance rather than Euclidean distance. For two cells *i* and *j* with the same label *t*_*i*_ = *t*_*j*_, the edge weight is

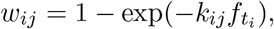

where *k*_*ij*_ is the spatial rank distance from cell *i* to cell *j*, and 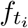 is the global fraction of cells assigned to label *t*_*i*_ within the analyzed section or dataset.

#### 1.1 Rank Distance Rather Than Euclidean Distance

Local cellular density can vary strongly across tissue. A fixed Euclidean radius would therefore correspond to very different numbers of nearby cells in dense and sparse regions. To reduce this dependence on local sampling density, we use rank distance, *k*_*ij*_, defined as the number of cells encountered when expanding outward from focal cell *i* until cell *j* is reached. Rank distance does not eliminate all density-dependent effects, but it makes the relevant spatial scale proportional to local cell count rather than physical distance alone.

#### 1.2 Null Motivation for the Edge Weight

The form of *w*_*ij*_ can be motivated by a simple complete-spatial-randomness null model. Let 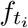 be the global fraction of cells of type *t*_*i*_. Under a perfectly mixed null, each cell encountered during radial expansion from *i* has probability 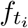 of also being type *t*_*i*_. Expanding to rank distance *k*_*ij*_ samples approximately *k*_*ij*_ cells, so the expected number of same-type cells encountered is

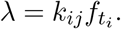

For large cell numbers and small local probabilities, the binomial count can be approximated by a Poisson random variable. Under this approximation, the probability of encountering zero same-type cells within rank distance *k*_*ij*_ is

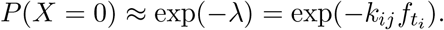

The complementary probability gives

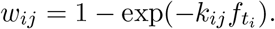

Thus, *w*_*ij*_ can be interpreted as an approximate cumulative waiting-time probability for encountering a same-type neighbor under a random mixing null. This frequency normalization gives rare and abundant labels more comparable edge-threshold meanings: a fixed threshold *P* corresponds to a similar null expectation for same-type encounter probability. This normalization does not make Map Information independent of class balance. Class imbalance still limits the amount of spatial information a label can contribute because very rare labels account for only a small fraction of cells, whereas overwhelmingly abundant labels are difficult to distinguish from the spatial background.

### 2 A Logistic Approximation for Non-Monotonicity

The empirical analyses show that Map Information is often maximized at an intermediate taxonomic granularity. This section gives an idealized explanation for that behavior. The goal is not to derive the exact score of the implemented graph, but to show how a non-monotonic relationship can arise from two opposing effects: increasing the number of labels can improve spatial specificity, but also increases the complexity of the categorical code.

#### 2.1 Assumptions of the Approximation

For this section, assume:

1. A map contains *N* cells assigned to *T* labels.
2. Labels are approximately balanced, so *H*_types_ ≈ log_2_ *T*.
3. The entropy trajectory *H*_*Z*_(*P*) can be approximated by a smooth sigmoidal collapse from *H*_cells_ = log_2_ *N* to *H*_types_ = log_2_ *T*.
4. The real map and the label-permuted null differ primarily in the location of this collapse along the proximity threshold *P*.

These assumptions are not required for computing Map Information. They are used only to illustrate the origin of the tradeoff.

#### 2.2 Approximate Percolation Entropy Curve

Let *F* (*P*; *P*_*c*_, *δ*) be a normalized sigmoid satisfying *F* (0; *P*_*c*_, *δ*) = 0 and *F* (1; *P*_*c*_, *δ*) = 1:

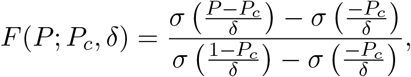

where

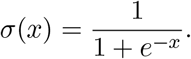

We approximate the zone entropy trajectory as

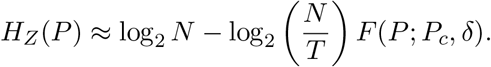

Here, *P*_*c*_ is an effective collapse point and *δ* controls the width of the transition. This is a phenomenological approximation to the percolation curve, not a claim that empirical percolation curves are exactly logistic.

#### 2.3 Approximate Expression for Map Information

Map Information is defined as

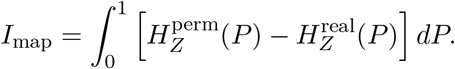

Substituting the logistic approximation gives

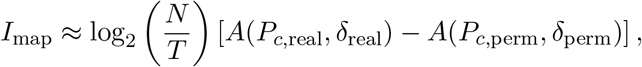

where

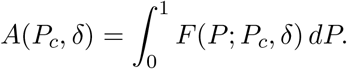

Because *F* (*P*) increases from 0 to 1, a real map that collapses earlier than its permuted null has a larger value of *A* and therefore a positive Map Information score.

#### 2.4 Mechanism of Non-Monotonicity

This approximation separates two effects. The first is the coding range,

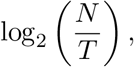

which decreases as the number of labels increases. This term represents the remaining addressable entropy that can be compressed by spatial organization. The second is the percolation advantage of the real map over its permuted null. Increasing *T* can improve this advantage if finer labels produce more spatially coherent domains and shift the real percolation curve leftward relative to the null.

The non-monotonic behavior arises when these effects compete. For small *T*, increasing the number of labels can reveal spatial structure that was hidden by overly coarse labels. In this regime, the percolation advantage can increase faster than the coding range decreases. For large *T*, further subdivision reduces the available coding range and can fragment otherwise coherent domains, causing the score to decline.

Two limiting cases are exact for the implemented definition:

- **Extreme under-clustering (***T* = 1): If all cells share one label, label permutation does not change the map. The real and permuted entropy curves are identical, so *I*_map_ = 0.
- **Extreme over-clustering (***T* = *N* ): If every cell has a unique label, no same-label spatial compression is possible. The type entropy equals the cellular address entropy, and *I*_map_ = 0.

Thus, any spatially structured intermediate taxonomy with positive Map Information must outperform both extremes. The logistic model illustrates why such an intermediate optimum is expected, although the exact location and height of the optimum depend on the candidate taxonomies and the empirical spatial graph.

### 3 Dependence of Map Information on Class Balance

Class balance affects Map Information because labels contribute to the entropy curve in proportion to the fraction of cells they describe. The rank-distance normalization in the edge weight makes proximity thresholds more comparable across rare and abundant labels, but it does not remove the consequences of class imbalance. Highly imbalanced labels have limited capacity to contribute to spatial compression.

#### 3.1 Additive Decomposition of Zone Entropy

Consider a binary map with *N* cells and class fractions *f* and 1 − *f*. Because the percolation graph only connects cells with the same label, the zone entropy can be decomposed into labelspecific contributions:

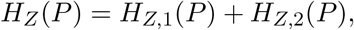

where

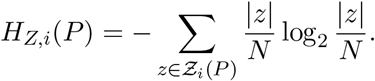

The corresponding Map Information score is

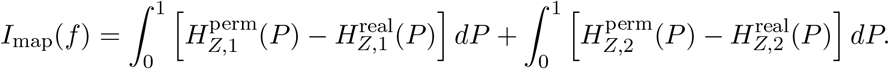

This decomposition is useful because it shows that each label contributes according to both its abundance and its spatial organization.

#### 3.2 Extreme Imbalance

Let Type 1 be rare, with *f* → 0, and Type 2 be the overwhelming majority. The rare class can only affect the total entropy through a vanishingly small fraction of the cells. Even if those rare cells are spatially organized, their total contribution to the entropy curve is limited by their abundance. At the same time, the majority class becomes nearly identical to the full tissue background, so its real and permuted spatial organization become increasingly similar. Therefore,

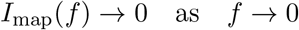

and by symmetry,

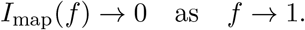

#### 3.3 Class Balance as an Entropy Envelope, Not a Strict Bound

The binary type entropy,

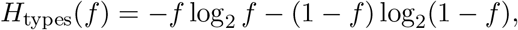

provides useful intuition for the abundance-dependent envelope of Map Information: balanced labels provide more opportunity for spatial compression than extremely imbalanced labels.

However, *H*_types_(*f*) should not be interpreted as a strict upper bound on the integrated Map Information score. Map Information integrates an entropy difference across proximity thresholds, and this integral can exceed the categorical entropy measured at a single endpoint.

The appropriate conclusion is therefore qualitative but important: class imbalance reduces the attainable contribution of a label to the score, whereas balanced and spatially coherent labels have greater potential to produce large Map Information. This is why Map Information favors taxonomies that use labels efficiently, but it does not imply that all labels must be perfectly balanced or that the score is bounded by type entropy alone.

### 4 The Unconstrained Spatial Model

The empirical analyses optimize over candidate taxonomies that are constrained by molecular or compositional feature spaces. This constraint is central: a cell-type taxonomy should not merely divide tissue into spatially compact domains; it should define labels that are recoverable from biological measurements. To clarify the importance of this distinction, this section considers an idealized *unconstrained* spatial model.

Here, “unconstrained” means that labels may be assigned directly from spatial position, without requiring support from molecular features, cell states, cell-type markers, or local cellular composition. This model is not intended to describe biological cell types. Instead, it defines a spatial compression reference case: what Map Information would favor if labels were allowed to optimize spatial compressibility alone. Comparing this reference case with feature-constrained taxonomies clarifies the central biological problem addressed by Map Information: finding spatially informative maps that remain grounded in measurable biological features.

#### 4.1 Assumptions of the Model

For analytical tractability, assume:

1. The tissue is represented by an idealized two-dimensional domain containing *N* cells.
2. The *N* cells are partitioned into *T* labels.
3. Labels are balanced, so each label contains *N/T* cells and *H*_types_ = log_2_ *T*.
4. In the real unconstrained map, each label occupies a compact spatial domain.
5. The entropy curves are approximated as step-like collapses rather than full empirical percolation trajectories.
6. The constants introduced below are geometry- and graph-dependent effective constants, not universal constants of the implemented metric.

The square-lattice language used below should be read only as a convenient reference geometry for scaling intuition. The implemented graph in the empirical analyses is a rank-distance, same-label graph, not a physical nearest-neighbor square lattice.

#### 4.2 The Real Map: Compact Domains

In the unconstrained optimum, a spatially efficient partition would place cells of the same label into compact domains. For such a map, same-label cells are close in rank distance and the entropy curve collapses early. Let *c* denote an effective local connectivity constant: the rank-distance scale, expressed in local-neighbor units, required for a compact same-label domain to become macroscopically connected under the approximate step-function model. This constant depends on geometry, graph construction, and finite-size effects. It is expected to be of order one for compact domains, but it is not fixed universally.

Using the implemented edge-weight form with balanced labels, *f*_*t*_ = 1*/T*,

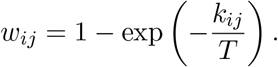

If compact-domain connectivity requires rank distance *k*_*ij*_ ≈ *c*, then the effective threshold for real-map collapse is

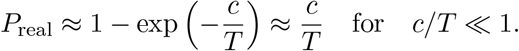

Approximating the real entropy curve as a step from log_2_ *N* to log_2_ *T* at this threshold gives

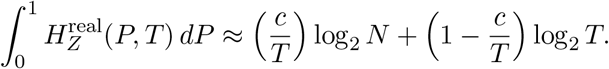

This approximation captures the intuition that compact domains can collapse early, but still pay a type-entropy cost at large *P*.

#### 4.3 The Permuted Map: Randomly Distributed Labels

In the label-permuted null, the *N/T* cells of each label are distributed randomly across the tissue. Under this idealized random-mixing model, the expected rank distance to a fixed-order same-label neighbor scales approximately as *T*, because the same-label density is 1*/T*. If the *m*-th same-label neighbor occurs at rank distance

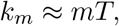

then the edge weight becomes

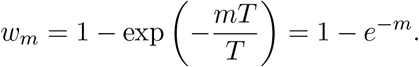

This cancellation shows why the frequency-normalized rank-distance graph makes the null percolation behavior approximately comparable across different numbers of balanced labels. The claim is not that the empirical null curve is exactly invariant to *T*, but that the leading dependence on label abundance is reduced by the normalization.

Let *A* denote the effective integrated fraction of the entropy range that remains above the final type entropy in the permuted null curve. Equivalently, *A* summarizes the area under the null entropy trajectory in the same step-function approximation. Its value depends on the reference geometry, graph construction, finite-size behavior, and approximation scheme. It should therefore be treated as a model parameter, not a universal constant. Under this approximation,

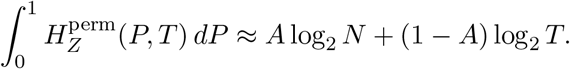

#### 4.4 Approximate Scaling of the Unconstrained Optimum

Combining the approximate real and permuted areas gives

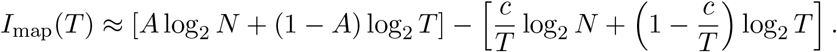

Rearranging,

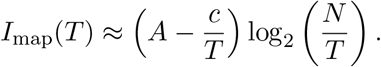

This expression illustrates the same tradeoff as in the empirical analyses. Increasing *T* improves the compact-domain advantage by reducing the real-map collapse threshold, but it also reduces the remaining addressable entropy log_2_(*N/T*).

Treating *T* as continuous within this idealized model, the stationary point satisfies

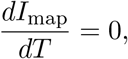

which gives

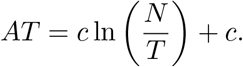

Equivalently,

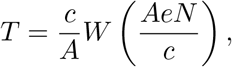

where *W* is the Lambert *W* function. For large *N*, this grows approximately logarithmically,

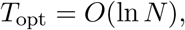

up to smaller corrections such as ln ln *N*. This scaling is a property of the idealized unconstrained model under the assumptions listed above. It should not be interpreted as a general law for biological cell-type number, nor as a prediction for the exact optimum in the empirical rank-distance graph. Its purpose is to show that even when labels are allowed to optimize spatial compression directly, Map Information favors an intermediate granularity rather than unlimited subdivision.

#### 4.5 Interpretation

The unconstrained model provides a useful reference because it separates two questions that are otherwise easy to conflate. The first is a spatial question: how compressible is the tissue if labels can be assigned directly from position? The second is a biological question: how much of that compressible spatial structure can be captured by labels derived from molecular or compositional features? The empirical analyses address the second question. The unconstrained model clarifies why the molecular constraint matters: without it, the optimal labels would be spatial domains, not necessarily biologically meaningful cell types.

### 5 Behavior Under Simple Misclassification Noise

This section considers a simplified noise model to illustrate how random misclassification can affect Map Information. The calculation is not a general proof of robustness. It assumes independent, spatially uncorrelated, symmetric label noise in an idealized compact-domain model. Real spatial transcriptomics errors may be correlated across space, enriched at boundaries, dependent on cell type, or biased toward specific confusions. Such errors can alter the percolation curves in ways not captured by the first-order calculation below.

#### 5.1 Assumptions of the Noise Model

We return to the unconstrained compact-domain model from Section 4. Assume that cells of type *t* occupy a dense spatial domain in the noise-free map. A fraction *ϵ* of cells in this domain are randomly assigned to incorrect labels, reducing the local density of correctly labeled type-*t* cells to

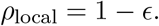

Assume further that the noise is symmetric across labels, so the global label fraction remains approximately unchanged:

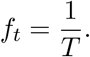

Under these assumptions, the label-permuted null is only weakly affected by the pre-existing random misclassifications, because the null already randomizes label positions. The dominant first-order effect is therefore modeled as a shift in the real-map percolation curve.

#### 5.2 Rank-Distance Expansion

In the noise-free compact domain, the effective rank distance required for local connectivity was approximated as

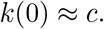

With random misclassification, correctly labeled cells are diluted by a factor 1 −*ϵ*. To encounter the same expected number of correctly labeled neighbors, the search must expand over a larger rank-distance neighborhood:

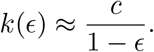

Substituting this into the edge-weight expression gives

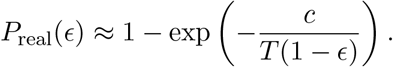

For *c/T* ≪ 1,

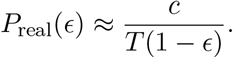

For small noise levels, *ϵ* ≪ 1,

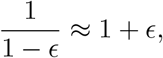

so

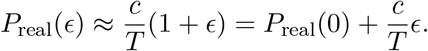

Thus, in this simplified model, independent random misclassification shifts the real-map collapse threshold to the right by approximately

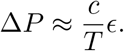

#### 5.3 Approximate First-Order Penalty

Using the same step-function entropy approximation as above, the penalty to Map Information is the threshold shift multiplied by the entropy range of the collapse:

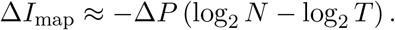

Therefore,

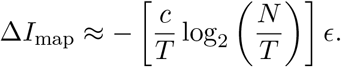

The resulting first-order approximation is

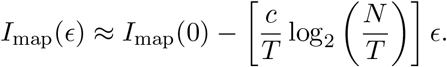

This calculation suggests that, under independent symmetric label noise, Map Information is expected to decrease gradually rather than collapse abruptly. The result should be interpreted as a local, first-order intuition for one simple noise regime. Correlated segmentation errors, boundary-localized errors, systematic cell-type confusion, or label-specific error rates could produce different effects and should be evaluated empirically for specific datasets.

---

**Supplementary Figure 1.**
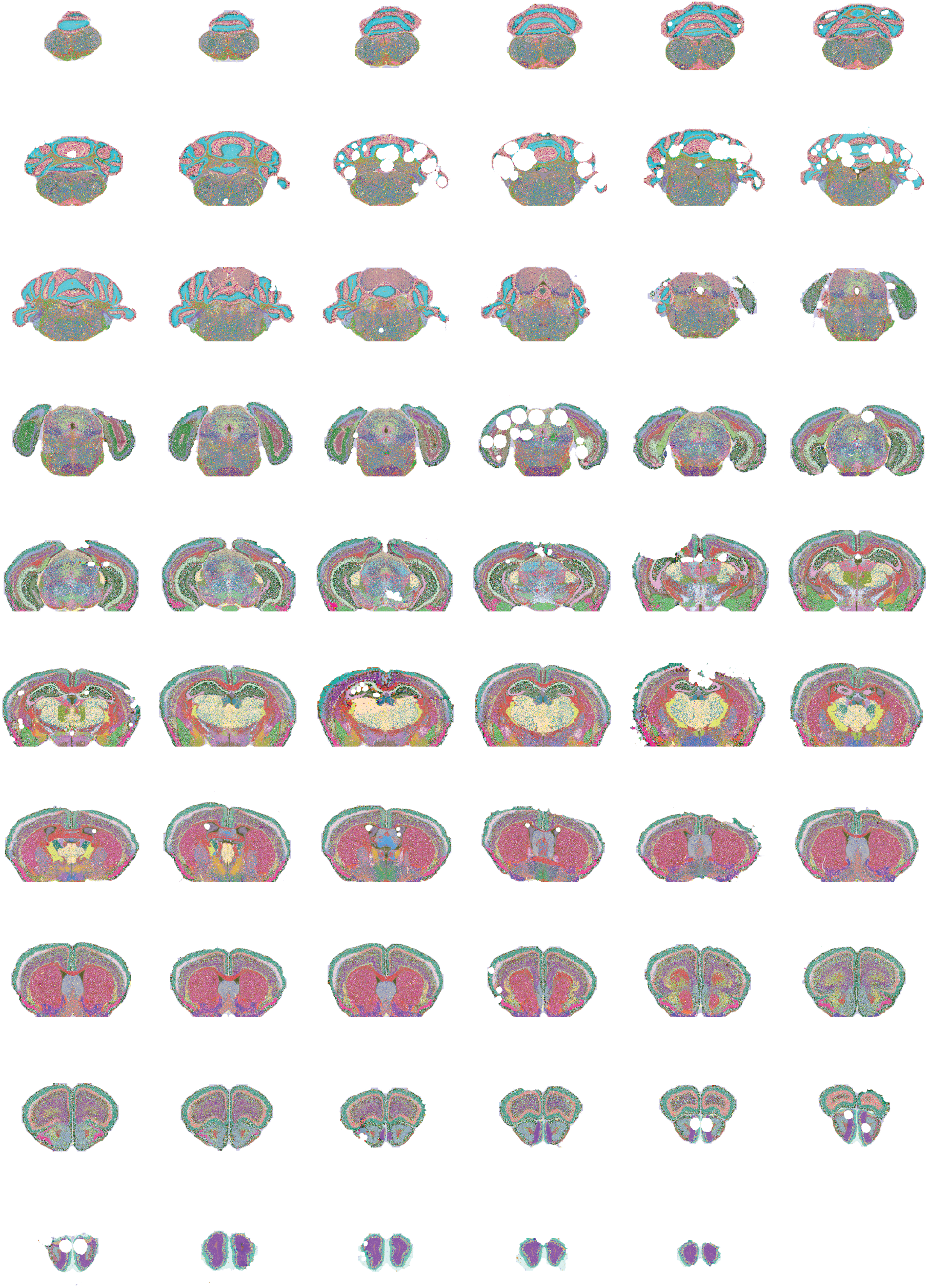
Optimized cell-type taxonomy across the entire mouse brain. Coronal sections are Shown in anterior-to-posterior order and colored by optimized cell-type identity.

**Supplementary Figure 2.**
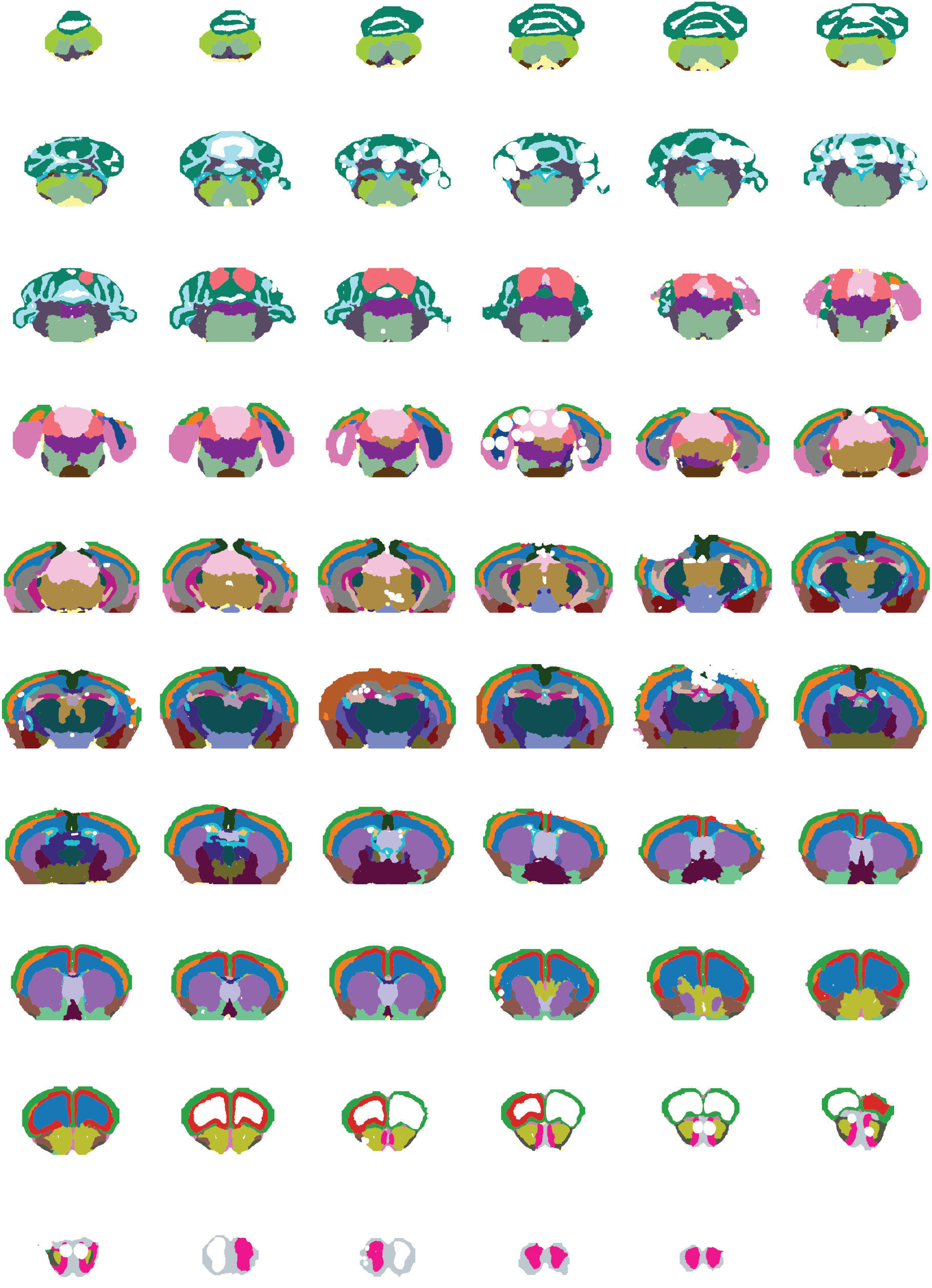
Optimized region taxonomy across the entire mouse brain. Coronal sections are shown in anterior-to-posterior order and colored by optimized region-type identity.

## References

[1] Lewis Carroll. Sylvie and Bruno Concluded. Macmillan and Co., 1893.

[2] Jorge Luis Borges. Del rigor en la ciencia. Los Anales de Buenos Aires, March 1946.

[3] Umberto Eco. How to Travel with a Salmon & Other Essays. Harcourt, 1994.

[4] Lewis Carroll. The Hunting of the Snark. Macmillan and Co., 1876.

[5] Vivien Marx. Method of the year 2020: Spatially resolved transcriptomics. Nature Methods, 18:9–14, 2021.

[6] William Bialek, Ilya Nemenman, and Naftali Tishby. Predictability, complexity, and learning. Neural Computation, 13(11):2409–2463, 2001.

[7] Raymond Cheong, Alexander Rhee, Carla J. Wang, Ilya Nemenman, and Andre Levchenko. Information transduction capacity of noisy biochemical signaling networks. Science, 334(6054):354–358, 2011.

[8] Jafar Selimkhanov, Brian Taylor, Junfeng Yao, Andrey Pilko, John G. Albeck, Alexander Hoffmann, Lev S. Tsimring, and Roy Wollman. Accurate information transmission through dynamic biochemical signaling networks. Science, 346(6215):1370–1373, 2014.

[9] Ryan Suderman, John A Bachman, Adam Smith, Peter K Sorger, and Eric J Deeds. Fundamental trade-offs between information flow in single cells and cellular populations. Proceedings of the National Academy of Sciences, 114(22):5755–5760, 2017.

[10] Evan Maltz and Roy Wollman. Quantifying the phenotypic information in mrna abundance. Molecular Systems Biology, 18:e11001, 2022.

[11] Julien O. Dubuis, Gasper Tkacik, Eric F. Wieschaus, Thomas Gregor, and William Bialek. Positional information, in bits. Proceedings of the National Academy of Sciences of the United States of America, 110(41):16301–16308, 2013.

[12] Vladimir Y. Kiselev, Tallulah S. Andrews, and Martin Hemberg. Challenges in unsupervised clustering of single-cell rna-seq data. Nature Reviews Genetics, 20(5):273–282, 2019.

[13] Detlev Arendt, Jacob M. Musser, Clare V. H. Baker, Aviv Bergman, Connie Cepko, Douglas H. Erwin, Mihaela Pavlicev, Gerhard Schlosser, Stefanie Widder, Manfred D. Laubichler, and Günter P. Wagner. The origin and evolution of cell types. Nature Reviews Genetics, 17(12):744–757, 2016.

[14] Naomi Moris, Cristina Pina, and Alfonso M. Arias. Transition states and cell fate decisions in epigenetic landscapes. Nature Reviews Genetics, 17(11):693–703, 2016.

[15] Zizhen Yao, Hanqing Liu, Fangming Xie, Stephan Fischer, Ricky S. Adkins, Andrew I. Aldridge, Seth A. Ament, Anna Bartlett, M. Margarita Behrens, Koen Van den Berge, et al. A transcriptomic and epigenomic cell atlas of the mouse primary motor cortex. Nature, 598(7879):103–110, 2021.

[16] Joseph M. Rich, Lambda Moses, Pétur Helgi Einarsson, Kayla Jackson, Laura Luebbert, A. Sina Booeshaghi, Sindri Antonsson, Delaney K. Sullivan, Nicolas Bray, Páll Melsted, and Lior Pachter. The impact of package selection and versioning on single-cell rna-seq analysis. Cell Systems, page 101560, 2026. In press.

[17] Shixiong Zhang, Xiangtao Li, Jiecong Lin, Qin Lin, Jie Chen, and Ka-Chun Wong. Review of single-cell rna-seq data clustering for cell-type identification and characterization. RNA, 29(5):517–530, 2023.

[18] Lingxi Yu, Yiran Cao, Rong Tang, Xuegong Wang, and Wencai Wang. Benchmarking clustering algorithms on estimating the number of cell types from single-cell rna-sequencing data. Genome Biology, 23(1):49, 2022.

[19] Breanne Sparta, Timothy Hamilton, Serena Hughes, Gokul Natesan, and Eric J. Deeds. A lack of distinct cell identities in single-cell measurements: revisiting waddington’s landscape. bioRxiv, 2025.

[20] Kenneth D. Harris, Hannah Hochgerner, Nathan G. Skene, Lucia Magno, Laszlo Katona, Cecilia Bengtsson Gonzales, Peter Somogyi, Nicoletta Kessaris, and Sten Linnarsson. Classes and continua of hippocampal ca1 inhibitory neurons revealed by single-cell transcriptomics. PLOS Biology, 16(6):e2006387, 2018.

[21] Garrett Stanley, Ozgun Gokce, Robert C. Malenka, Thomas C. Südhof, and Stephen R. Quake. Continuous and discrete neuron types of the adult murine striatum. Neuron, 105(4):688–699.e8, 2020.

[22] Tara Chari and Lior Pachter. The specious art of single-cell genomics. PLOS Computational Biology, 19(8):e1011288, 2023.

[23] Shamus M. Cooley, Timothy Hamilton, Eric J. Deeds, and J. Christian J. Ray. A novel metric reveals previously unrecognized distortion in dimensionality reduction of scrna-seq data. bioRxiv, 2022.

[24] Fuchou Tang, Catalin Barbacioru, Yangzhou Wang, Ellen Nordman, Chuanchao Lee, Ning Xu, Xiaohui Wang, John Bodeau, Brian B. Tuch, Asim Siddiqui, Kai Lao, and M. Azim Surani. mrna-seq whole-transcriptome analysis of a single cell. Nature Methods, 6(5):377–382, 2009.

[25] Patrik L. Ståhl, Fredrik Salmén, Sanja Vickovic, Anna Lundmark, José F. Navarro, Jens Magnusson, Stefania Giacomello, Michaela Asp, Jakub O. Westholm, Mikael Huss, Annelie Mollbrink, Sten Linnarsson, Simone Codeluppi, Åke Borg, Fredrik Pontén, Paul I. Costea, Pelin Sahlén, Jan Mulder, Olaf Bergmann, Joakim Lundeberg, and Jonas Frisén. Visualization and analysis of gene expression in tissue sections by spatial transcriptomics. Science, 353(6294):78–82, 2016.

[26] Orit Rozenblatt-Rosen, Michael J. T. Stubbington, Aviv Regev, and Sarah A. Teichmann. The human cell atlas: from vision to reality. Nature, 550(7677):451–453, 2017.

[27] Zizhen Yao, Caleb T. J. van Velthoven, T. N. Nguyen, Jeff Goldy, Adriana E. Sedeno-Cortes, Farzaneh Baftizadeh, Darren Bertagnolli, Tim Casper, Katharyn Crichton, Shiella-Lyn Ding, et al. A high-resolution transcriptomic and spatial atlas of cell types in the whole mouse brain. Nature, 624(7991):317–332, 2023.

[28] Vipul Singhal, Nigel Chou, Joseph Lee, et al. Banksy unifies cell typing and tissue domain segmentation for scalable spatial omics data analysis. Nature Genetics, 56:431–441, 2024.

[29] Zhiyuan Yuan et al. Mender: fast and scalable tissue structure identification in spatial omics data. Nature Communications, 15:207, 2024.

[30] Zhiyuan Yuan, Fangyuan Zhao, Senlin Lin, et al. Benchmarking spatial clustering methods with spatially resolved transcriptomics data. Nature Methods, 21:660–672, 2024.

[31] Quanxin Wang, Song-Lin Ding, Yang Li, Josh Royall, David Feng, Paul Lesnar, Nina Graddis, Maryam Naeemi, Brian Facer, Allen Ho, Tim Dolbeare, Brent Blanchard, Nick Dee, Wayne Wakeman, Kellen E. Hirokawa, Aaron Szafer, Susan M. Sunkin, Seung W. Oh, Allan Bernard, Jeremy W. Phillips, Michael Hawrylycz, Christof Koch, Hongkui Zeng, Julie A. Harris, and Lydia Ng. The allen mouse brain common coordinate framework: A 3d reference atlas. Cell, 181(4):936–953.e20, 2020.

[32] Hongkui Zeng. What is a cell type and how to define it? Cell, 185(15):2739–2755, 2022.

[33] Hans Clevers, Susanne Rafelski, Michael Elowitz, Allon Klein, Jay Shendure, Oliver Stegle, Amos Tanay, Cole Trapnell, Ed Lein, Emma Lundberg, et al. What is your conceptual definition of “cell type” in the context of a mature organism? Cell Systems, 4(3):255–259, 2017.

[34] Kara L. McKinley, Diego Castillo-Azofeifa, and Ophir D. Klein. Tools and concepts for interrogating and defining cellular identity. Cell Stem Cell, 26(5):632–656, 2020.

[35] Silvia Domcke and Jay Shendure. A reference cell tree will serve science better than a reference cell atlas. Cell, 186(6):1103–1114, 2023.

[36] Monika Sester. Cartographic generalization. Journal of Spatial Information Science, Vol.2020(21):5–11, 2020.

[37] Paul A. David. Clio and the economics of qwerty. The American Economic Review, 75(2):332–337, 1985.

[38] Meng Zhang, Steven W. Eichhorn, Brian Zingg, Zizhen Yao, Keith Cotter, Hongkui Zeng, and Hong-Wei Dong. Spatially resolved cell atlas of the mouse brain by merfish. Nature, 598:137–143, 2021.

[39] Rafael Yuste, Michael Hawrylycz, Nadia Aalling, Argel Aguilar-Valles, Detlev Arendt, Rubén Armañanzas, Giorgio A. Ascoli, Concha Bielza, Vahid Bokharaie, Till B. Bergmann, Iryna Bystron, Marco Capogna, Yun Jeong Chang, Ann Clemens, Christiaan P. J. de Kock, Javier DeFelipe, Sandra E. Dos Santos, Kate Dunville, Dirk Feldmeyer, Richárd Fiáth, Gord Fishell, Alberto Foggetti, Xin Gao, Payam Ghaderi, Natalia A. Goriounova, Onur Güntürkün, Kosuke Hagihara, Vanessa J. Hall, Moritz Helmstaedter, Suzana Herculano-Houzel, Markus M. Hilscher, Hajime Hirase, Jens Hjerling-Leffler, Rebecca Hodge, Z. Josh Huang, Rafiq Huda, Konstantin Khodosevich, Ole Kiehn, Christof Koch, Eric S. Kuebler, Malte Kühnemund, Pedro Larrañaga, Boudewijn Lelieveldt, Emily L. Louth, Jan H. Lui, Huib D. Mansvelder, Oscar Marin, Julio Martinez-Trujillo, Zoltán Molnár, A. N. Mohapatra, Helena Munguba, Maiken Nedergaard, Pavel Nëmec, Nitzan Ofer, Ulrich G. Pfisterer, Susana Pontes, William Redmond, Jean Rossier, Joshua R. Sanes, Richard H. Scheuermann, Esther Serrano-Saiz, Jochen F. Staiger, Peter Somogyi, Gábor Tamás, Andreas S. Tolias, Maria A. Tosches, Marta Trejo García, Claudia Wozny, Tobias V. Wuttke, Yong Liu, Jing Yuan, Hongkui Zeng, and Ed S. Lein. A community-based transcriptomics classification and nomenclature of neocortical cell types. Nature Neuroscience, 23(12):1456–1468, 2020.

[40] Wei Dong, Charikar Moses, and Kai Li. Efficient k-nearest neighbor graph construction for generic similarity measures. In Proceedings of the 20th international conference on World wide web, pages 577–586. ACM, 2011.

[41] Vincent A Traag, Ludo Waltman, and Nees Jan Van Eck. From louvain to leiden: guaranteeing well-connected communities. Scientific reports, 9(1):5233, 2019.

[42] Jacob H. Levine, Erin F. Simonds, Sean C. Bendall, Kara L. Davis, El-ad D. Amir, Michelle D. Tadmor, Oren Litvin, Harris G. Fienberg, Astraea Jager, Eli R. Zunder, Rachel Finck, Amanda L. Gedman, Ina Radtke, James R. Downing, Dana Pe’er, and Garry P. Nolan. Data-driven phenotypic dissection of AML reveals progenitor-like cells that correlate with prognosis. Cell, 162(1):184–197, 2015.

[43] Brian C Ross. Mutual information between discrete and continuous data sets. PloS one, 9(2):e87357, 2014.

